# droplet-Tn-Seq combines microfluidics with Tn-Seq identifying complex single-cell phenotypes

**DOI:** 10.1101/391045

**Authors:** Derek Thibault, Stephen Wood, Paul Jensen, Tim van Opijnen

## Abstract

While Tn-Seq is a powerful tool to determine genome-wide bacterial fitness in high-throughput, culturing transposon-mutant libraries in pools can mask community or other complex single-cell phenotypes. droplet-Tn-seq solves that problem by microfluidics facilitated encapsulation of individual transposon mutants into liquid-in-oil droplets, thereby enabling isolated growth, free from the influence of the population. Importantly, all advantages of Tn-Seq are conserved, while reducing costs and greatly extending its applicability.

## Main text

Transposon insertion sequencing (Tn-Seq) has become the gold standard to determine in high-throughput and genome-wide a gene's quantitative contribution to fitness under a specific growth condition (van Opijnen et al. 2009). It has been successfully applied to bacteria, yeast and eukaryotes and has enabled the discovery of new biology including gene function, non-coding RNAs and host-factors affecting disease susceptibility (van Opijnen & Camilli 2013). One of the biggest strengths of Tn-Seq is the ability to screen hundreds of thousands of mutants in a single experiment. However, growing mutants *en masse,* i.e. in a pool, can mask the fitness of certain mutants. For instance, secreted factors that break down complex glycans into smaller units for energy utilization, can be viewed as community factors since mutants that do not produce these enzymes can 'cheat' and reap the carbon-source benefits. Moreover, additional mechanisms including frequency dependent selection, bet-hedging and division of labor can retain mutants with a relatively low individual fitness in a population, which are all missed by Tn-Seq (Veening et al. 2008; Sæther & Engen 2015).

In order to obtain a comprehensive understanding of a complex population it is thus important to consider the fitness of each individual in isolation as well as in the context of the population. To achieve this we developed droplet-Tn-Seq (dTn-Seq), by combining Tn-Seq with microfluidics. In dTn-Seq a microfluidic device enables encapsulation of millions of single bacterial cells in micron-sized droplets in which bacteria are cultured. Each transposon mutant thus starts off in a complex pool of mutants, is then separated and cultured in isolation, and finally cells are pooled back together. Before encapsulation and after pooling, genomic DNA is isolated for sample preparation and the change in frequency of each mutant over the course of the experiment is determined through massively parallel sequencing, which is used to calculate individual growth rates (van Opijnen et al. 2009). Therefore, through strategic isolation and pooling, dTn-Seq enables the establishment of single cell behavior in a genome-wide and high-throughput fashion (Fig. 1a). Moreover, we show that besides the ability to resolve complex single-cell behavior, droplets have many more advantages and applications including a drastic reduction in culture media volume (and possible expensive compounds), and analyses of interactions between bacteria and/or host-cells.

**Figure 1:**
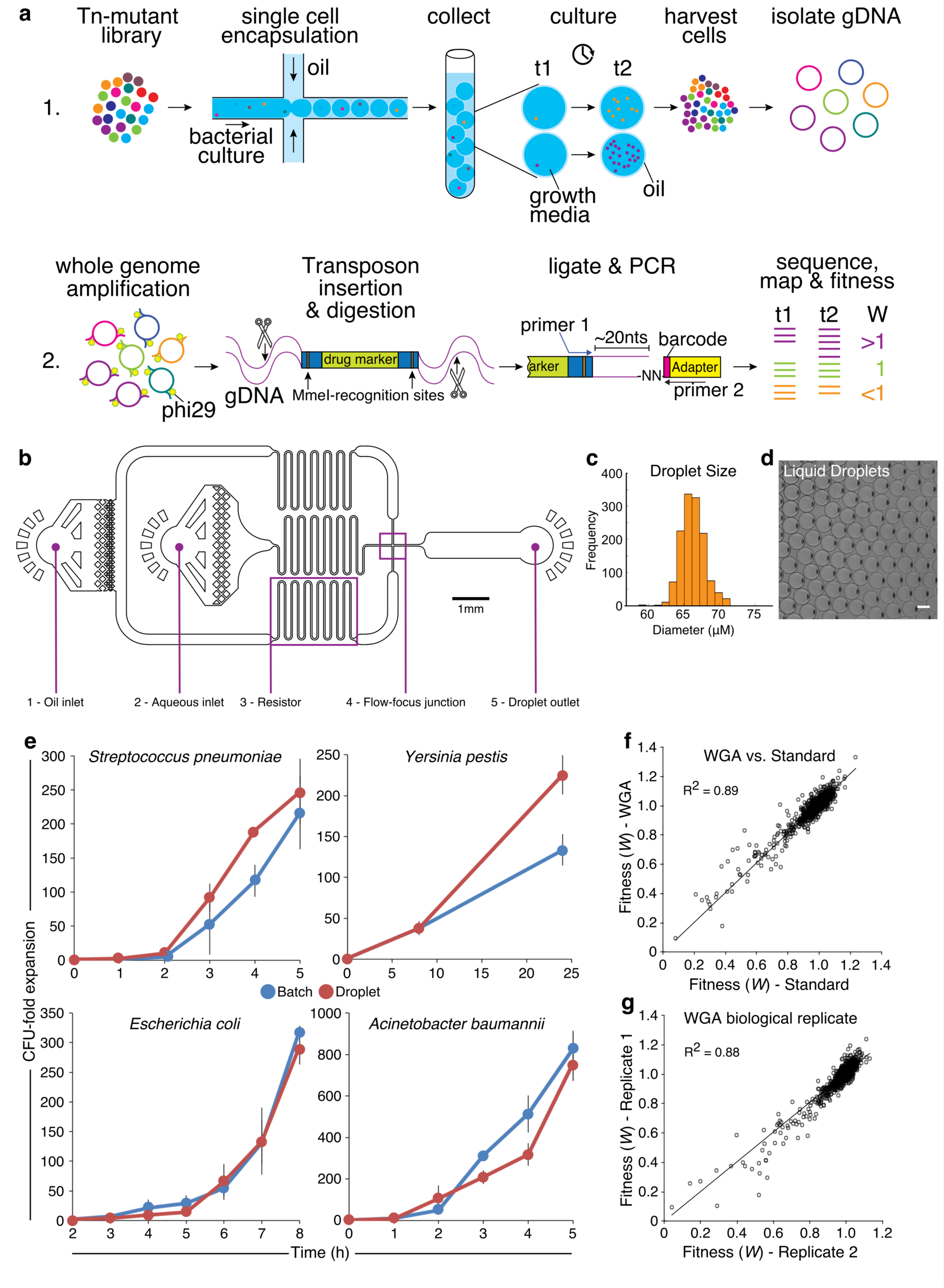
Droplet Tn-Seq overview and characterization. **(a)** A microfluidic device encapsulates single bacterial cells into liquid-in-oil droplets. Bacteria are allowed to expand within droplets, gDNA is isolated at the start of the experiment (t1) and after expansion (t2) and is amplified with phi29. Importantly, while expansion for each transposon mutant takes place in isolation, gDNA is isolated from the pooled population, enabling screening of all mutants simultaneously. gDNA is digested with MmeI, an adapter is ligated, a ~180bp fragment is produced which contains approximately 16 nucleotides of bacterial genomic DNA, defining the transposon insertion location, followed by Illumina sequencing. Reads are demultiplexed (based on the barcode in the adapter and a potential second barcode in primer 1), mapped to the genome, and fitness is calculated for each defined region. (**b**) Syringe pumps and tubing are used to deliver surfactant in a fluorinated carrier oil to the Oil-inlet (1) and culture media containing cells to the Aqueous inlet (2). Resistors (3) reduce fluctuation in liquid flow rates. Oil separates the continuous flow of the cell culture into monodisperse droplets at the Flow-focus junction (4). Droplets exit the device through the Droplet-outlet (5) and are collected. (**c**) Depending on the size of the channels, and oil and aqueous phase flow rates, uniformly sized droplets can be formed ensuring each cell has the same expansion potential. With a channel size of 40 µM ~67 µM diameter droplets are created. (**d**) Liquid droplets in carrier oil. Scale bar, 50μM. (**e**) Both Gram-positive and negative bacteria expand robustly in liquid droplets, in a similar fashion to batch culture (i.e. a 5ml culture). Depending on the experiment the amount of gDNA may be limited, which can be resolved by whole genome amplification (WGA), which introduces no bias compared to a standard Tn-Seq library prep (**f**) and is reproducible (**g**).

To enable encapsulation of single cells into a single droplet, microfluidic devices were designed and built in-house (Fig. 1b, Supplementary File 1), which depending on oil and aqueous phase flow rates generate droplets with a diameter of ~67 µM at a rate of ~5×10^4^ droplets/minute. Droplets are composed of an outer oil-surfactant layer and are filled with growth media (~157pL; Fig. 1c, d). They enable robust bacterial growth for Gram-negative and positive bacteria alike and support overall growth dynamics comparable to bacteria grown in batch culture (a large e.g. 5ml culture; Fig. 1e). Pooled transposon libraries, where each bacterial cell contains a single transposon insertion, are separated as single cells by encapsulation and cultured inside the droplet for 5-8 generations. With (d)Tn-Seq, massively parallel sequencing is used to determine the exact location of a transposon insertion and by sequencing two time points (i.e. at the start and end of the experiment) the frequency of each mutant can be accurately quantified which is used to calculate the transposon's impact on the growth rate (van Opijnen et al. 2009, van Opijnen & Camilli 2013). Due to the small volume of the droplet the amount of genomic DNA obtained from a dTn-Seq experiment is not sufficient for sample preparation. To overcome this a whole genome amplification (WGA) step, mediated by phi29 is introduced. WGA conditions were optimized so that when dTn-Seq library preparation is compared against the standard Tn-Seq approach no bias is observed (R^2^=0.89; Fig. 1f) and reproducibility is very high (R^2^=0.88; Fig. 1g).

To determine the functionality of dTn-Seq, transposon insertion libraries of *Streptococcus pneumoniae* were grown in batch-culture as a pooled population ('standard' Tn-Seq) and encapsulated as single cells (dTn-Seq) in growth media with either glucose or the complex glycan alpha-1-acid glycoprotein (AGP) as the major carbon source. Moreover, by adding low-melting temperature agarose to growth media with glucose, droplets with a 1% agarose density were generated to assess how a solid environment that provides structural support affects single cell growth. For each gene in each condition, fitness (i.e. the growth rate) was calculated and compared between pooled-batch and droplet conditions. Overall, 2-5% of genes from a variety of categories, including metabolism, transport, regulation and cell wall integrity have a significantly different fitness (Supplementary Fig. 1), indicating that population structure, i.e. the droplet environment, can significantly affect clonal fitness.

To validate dTn-Seq a total of 7 genes were chosen from the three environments (Supplementary Fig. 1; Supplementary Tables 1-5). In the simplest environment with glucose as the carbon source *lytB* (SPT_1238) has no effect on fitness, however when grown in droplets the mutant has a severe growth defect (Fig. 2a, b; Supplementary Fig. 1; Supplementary Table 1). This defect seems to be due to the small droplet environment, since poor growth is masked when the mutant is grown by itself in batch culture (5ml; Fig. 2b). LytB is part of the lytic cycle of *S. pneumoniae* and is involved in cell-chain shortening (García et al. 1999). Indeed, chain-length of *lytB* is significantly longer then the wt when grown in batch, however when the mutant is grown in droplets, chain-lengths are shortened and indistinguishable from wt (Fig. 2a). Recently longer cell chains were associated with rapid local induction of competence (Domenech et al. 2018), which we hypothesized could be further enhanced in the micro-droplet environment. Gene-expression of a set of competence genes was compared between wt and *lytB* grown in batch and in droplets. As posited, competence genes of *lytB* cultured in droplets are highly upregulated (Fig. 2c). Importantly, competence also induces the autolysin *cbpD* as well as the immunity gene *comM*. In *lytB*-droplets *comM* is upregulated ~8-fold, while *cbpD* is upregulated ~28- fold. This means that, especially in a confined space, LytB is extremely important in limiting a local hypercompetent phenotype, which when deleted triggers autolysis and fratricide and reduces fitness.

**Figure 2:**
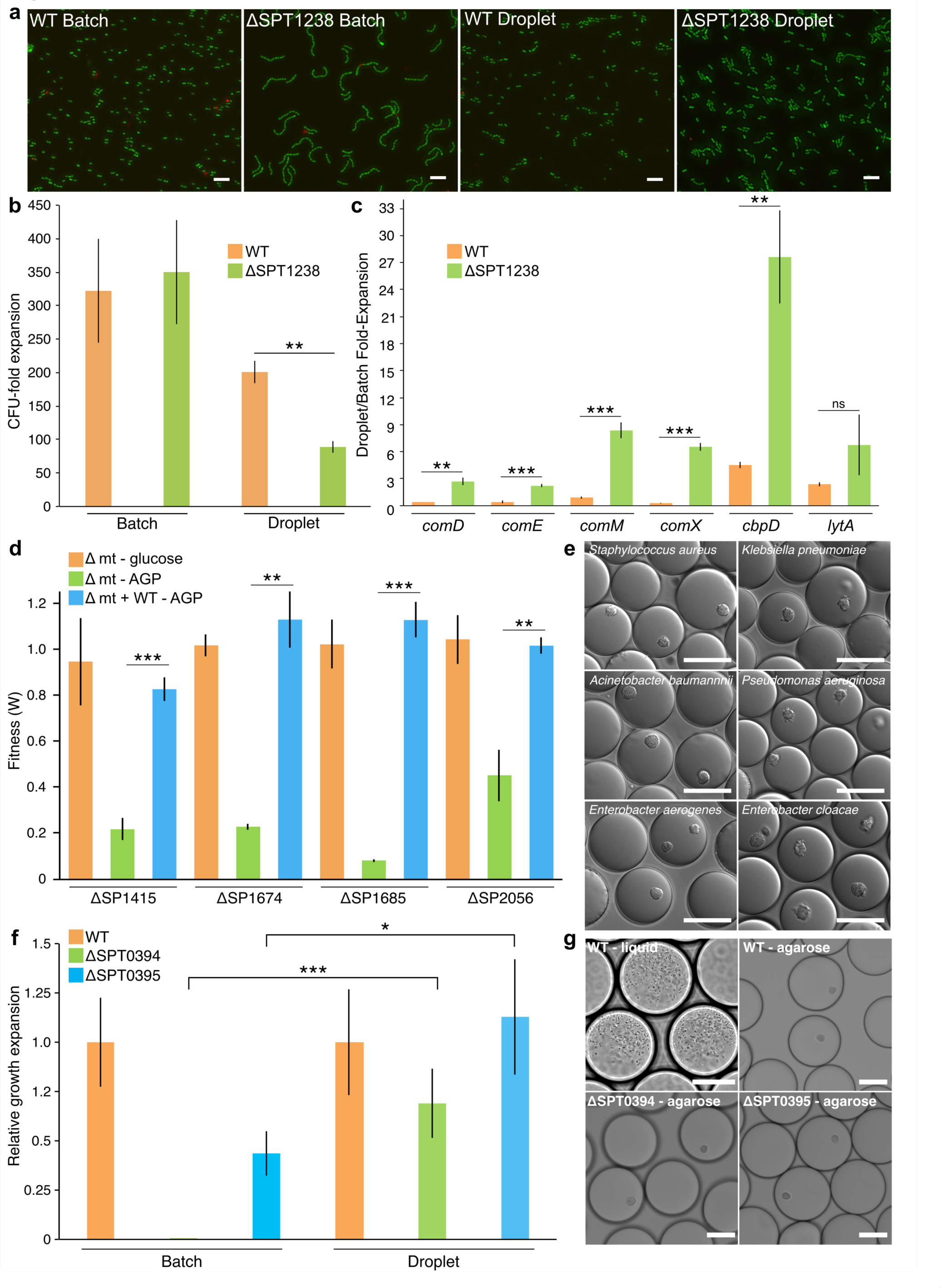
dTn-Seq validation. (**a**) Wt has a shorter chain length then *lytB* in batch culture, but in droplets chain lengths are similar. (**b**) Wt and *lytB* expand and grow at a similar rate in batch-culture, however, in droplets *lytB* expands less then wt. Shorter chain lengths and less expansion of *lytB* in droplets could either be caused by slower growth or a higher death rate. (**c**) Gene expression analyses shows significant upregulation of competence genes *comD, E, M,* and *X*, with the autolysin *cbpD* being upregulated ~28-fold in droplets, indicating that fratricide and thus an increased death rate is limiting growth of *lytB* in droplets. Moreover, this indicates a new role for LytB, which is to suppress local hyper-competence. (**d**) While deletion mutants of SP_1415, SP_1674, SP_1685, and SP_2056 have no defect when grown independently in medium with glucose as the carbon source (orange bars), they hardly grow when glucose is replaced by AGP (green bars). Importantly, this growth defect in AGP can be resolved by adding wt to the culture (blue bars), indicating that wt is providing 'community support'. (**e**) Agarose droplets can be generated by adding low melting agarose to growth media, which provides structural support and results in bacteria (Gram-negative and positive alike) growing in microcolonies. Scale bars 50μM. (**f, g**) Two capsule mutants (SPT_0394 and SPT_0395) that grow (very) slow in liquid batch culture and liquid droplets, expand robustly in agarose droplets. *p<0.05, **p<0.005, ***p<0.0005 in a *student t-test*.

We next compared transposon libraries grown in culture medium with AGP as the major carbon source. AGP is a highly glycosylated protein found in serum with covalently linked carbohydrate side chains composed of linked monosaccharides such as mannose, galactose, N-acetylglucosamine (GlcNAc/GlcN), and sialic acid. Most bacteria, including *S. pneumoniae,* are unable to take up such large structures and depend on monosaccharides being liberated, for instance by secreted enzymes (King 2010). Four genes (SP_1415/*nagB*, SP_1674, SP_1685/*nanE*, SP_2056/*nagA*) with a severe growth defect in AGP-droplets and no defect in batch culture were validated. Each gene is dispensable when grown by itself in media with glucose but is highly important for growth when AGP is the carbon source (Fig. 2d; Supplementary Fig. 1; Supplementary Table 4). Importantly, when a deletion mutant of either gene is grown in the presence of the wildtype the growth defect in AGP is masked (Fig. 2d), indicating that the wildtype is providing community support and compensating the mutant's fitness. While none of the four genes have previously been shown to be influenced by the community, each gene is associated with either regulating, releasing and/or processing AGP-linked monosaccharides. Specifically, SP_1415/*nagB* and SP_2056/*nagA* have been shown in other species to be involved in processing GlcN and GlcNAc (Moye et al. 2014), SP_1674 is a predicted transcriptional activator of a regulon containing *nanA* and *nanB* which have been show to release sialic acid from complex glycan structures (King 2010), and SP_1685/*nanE* is a putative lipoprotein anchored to the membrane and important for sialic acid utilization (Pélissier et al. 2014). These data show that dTn-Seq is highly sensitive in identifying genes and processes that can be shared amongst bacteria and enable 'cheating', which are missed with Tn-Seq.

Lastly, two capsule genes (SPT_0394/*cpsC*, SPT<u>_</u>0395/*cpsD*) were validated that are very important for growth under standard conditions (e.g. liquid media), but whose fitness is largely compensated by the addition of 1% agarose (Fig. 2f; Supplementary Fig. 1; Supplementary Table 2). Like liquid droplets, agarose droplets are monodisperse and have a similar volume (Supplementary Fig. 2). Indeed, when the deletion mutants are cultured in liquid droplets or in batch, SPT_0394 hardly grows, while SPT_0395 grows slower than the wt, reflecting their fitness (Fig. 2f). In contrast, the 1% agarose environment allows both mutants to expand robustly, and form microcolonies similar to wt (Fig. 2f,g). Microcolony formation is important for bacterial survival in host-tissue, and for instance makes *Pseudomonas aeruginosa* less sensitive to antimicrobials (Lam et al. 1980; Worlitzsch et al. 2002; Sriramulu et al. 2005). Microcolonies and biofilms are both formed by clusters of bacteria and thus dTn-Seq could provide a proxy to uncover genes that are important under such circumstances. Importantly, noncapsular *S. pneumoniae* strains are often better at biofilm formation (Domenech et al. 2012), which is suggestive for the improved performance of the capsule mutants in agarose.

To conclude, dTn-Seq is a valuable extension of Tn-Seq that is able to uncover novel single-cell phenotypes associated with microenvironments, community factors, and solid environments that are masked by Tn-Seq. dTn-Seq is applicable to practically any bacterium and any variation of Tn-Seq (e.g. IN-Seq, TraDIS, HITS, Bar-Seq). Importantly, we have only shown a limited number of environments but there are many other possibilities. For instance, we have successfully used dTn-Seq in combination with antibiotics, in a screen for siderophores, and to determine interactions between bacteria and host-cells. Moreover, agarose droplets are sortable via FACS and droplets are easily imaged. Lastly, the small droplet environment reduces the amount of (expensive) compounds and chemicals needed to perform an experiment (e.g. AGP) thereby enabling genome-wide studies for a fraction of the cost.

## Methods

### Bacterial strains, growth and media

Sequencing and validation experiments were performed using *Streptococcus pneumoniae* TIGR4 (NCBI Reference Sequence: NC_003028.3), and Taiwan-19F (NC_012469.1). Other species used in the study were *Yersinia pestis* (KIM6, pCD1 negative and pgm negative), *Escherichia coli* (DH5-α), *Acinetobacter baumannii* (ATCC 17978), *Staphylococcus aureus* (RN1, NR-45904), *Klebsiella pneumoniae* (UHKPC57, NR-44357), *Pseudomonas aeruginosa* (PA14), *Enterobacter aerogenes* (NRRL B-115), and *Enterobacter cloacae* (NRRL B-412). Except for specific growth and selection experiments the *S. pneumoniae* strains were cultured statically in Todd Hewitt broth supplemented with yeast extract (THY) plus 5 µl/ml Oxyrase (Oxyrase, Inc.) and 150 U/ml catalase (Worthington Bio Corp LS001896), or on Sheep's blood agar plates at 37°C in a 5% CO_2_ atmosphere. *Y. pestis* was cultured in brain heart infusion media or on blood agar while all other strains were cultured in Luria-Bertani (LB) broth or on LB agar at 37°C. Unless otherwise noted cells were cultured to exponential phase before being washed in PBS and diluted down into the appropriate media.

### Transposon library construction and selection experiments

Library construction using the mariner transposon Magellan6 was performed as previously described (van Opijnen & Camilli 2013; van Opijnen & Camilli 2012; van Opijnen et al. 2009). The transposon lacks transcriptional terminators allowing for read-through transcription, and additionally has stop codons in all three frames in either orientation to prevent aberrant translational products. Six independent transposon libraries were produced for each experimental condition using *S. pneumoniae* strains TIGR4 or Taiwan-19F. Each transposon library consists of at least 10,000 total mutants. The environmental conditions for selection experiments included growth in semi-defined minimal media (SDMM) (van Opijnen & Camilli 2012) at pH 7.3 supplemented with 20 mM glucose, human alpha-1-acid glycoprotein (Sigma - G9885), and agarose (Lonza - Seaplaque, 50101). Every selection experiment was cultured at 37°C in a 5% CO_2_ atmosphere.

### Microfluidic device production

Microfluidic device masks were designed using AutoCad 2016 software (AutoDesk) (Supplemental File 1) and photomasks were ordered from CAD/Art Services, Inc. (Bandon, OR). The mold and final microfluidic chip fabrication was performed at the Integrated Sciences Cleanroom and Nanofabrication Facility at Boston College. A master mold was fabricated by coating a silicon wafer with SU-8 3025 (MicroChem) using a spin coater (Laurell) and set by baking at 95°C. The photomask was aligned with the silicon wafer and UV exposed followed by a post-exposure bake ramping from 65°C to 95°C over 4 minutes. The mold was developed using SU-8 developer (MicroChem) per the manufacturers guidelines and rinsed with isopropanol and dH2O followed by a hardening bake from 100C to 200C over 5 minutes. The PDMS chip was generated by mixing PDMS and curing agent (Dow Corning, Sylgard 184) in a 10:1 ratio and added to the mold, degassed with a vacuum, and polymerized at 65°C overnight. Polymerized PDMS was cut from the mold and a biopsy punch (0.75mm - Shoney Scientific) was used to create ports for tubing (PE-2 tubing - Intramedic). PDMS slabs were bonded to glass (Corning - 2947, 75×50mm) at the clean room by washing the glass with acetone and isopropanol in a sonicator bath while the PDMS was washed with isopropanol, followed by thorough drying with filtered nitrogen gas. The channel side of the PDMS slab and the glass slide were treated with plasma (400sccm flow; 400 watts; 45 sec) using a faraday barrel screen. Plasma treated surfaces were quickly brought into contact and pressed together and then placed at 65°C for 10 minutes to complete bonding.

### Droplet production and culturing of bacteria in liquid and agarose droplets

Before droplet production the device's aqueous channel was primed with Aquapel (Aquapel #47100) and then flushed with fluorinated oil (Novec 7500 oil; 3M #98-0212-2928-5) (Fig. 1b). Devices were used immediately or incubated overnight at 65°C, covered in scotch tape, and then stored in the dark for several weeks before use. A 1ml syringe (BD - 309628) was filled with 1.5% of PicoSurf-1 in Novec 7500 oil (Dolomite; 3200214) while another 1ml syringe was filled with cell culture and then both were hooked into syringe pumps (Cole-Parmer Instrument Co. - 00280QP). PE-2 tubing was used to transfer PicoSurf-1 oil to the 'oil inlet', cell culture to the 'aqueous inlet', and collect droplets from the 'droplet outlet' (Fig. 1b). Cells were diluted based on droplet size and according to a Poisson distribution with the goal of generating droplets that contained a single bacterial cell (Shapiro 2003). With our device a concentration of 2.1 × 10^6^ cells/ml encapsulated into ~157pL sized droplets will yield approximately 74% empty droplets, 22% with single cells, and 3% with two or more cells. The syringe pump rate for cell encapsulation was 400 μl hr^-1^ yielding ~1.4×10^6^ total droplets in 30 minutes. To generate agarose droplets the entire droplet production system was placed in a 37°C warm room. 1% Seaplaque agarose was added to growth media and then heated until dissolved. The agarose was then filtered (0.22μm) after which the cells were added to the agarose solution. After production, agarose droplets were gelled at 4°C for 10 min with occasional shaking, and then transferred to the incubation chamber. To produce growth curves small fractions of droplet culture were collected and broken open with 1H,1H,2H,2H-perfluoro-1-octanol (PFO; Sigma-Aldrich - 370533), which separated oil and aqueous culture phases. For liquid droplets the aqueous culture phase was immediately plated for live cell counts while the aqueous phase for agarose droplets was added to a dounce homogenizer to break up the agarose to release cells for live cell plating. CFU-fold expansion was calculated by dividing CFU counts at every time point by the initial CFU count at the beginning of each experiment. A *student's t-test* was used to determine if expansion between samples was significantly different (*p<0.05, **p<0.005, ***p<0.0005).

### DNA sample preparation, Illumina sequencing, and fitness calculation

Genomic DNA (gDNA) was extracted using the DNeasy Blood & Tissue Kit according to the manufacturer's guidelines for Gram-positive bacteria (Qiagen - 69506). DNA adapter barcodes were made by mixing an equal volume of primers ADBC-F and ADBC-R (Supplementary Table 6) at a concentration of 0.2 nM in buffer (10mM Tris-base, 50mM NaCl, 1mM EDTA, pH 8), followed by incubation at 95°C for 3 min, 60°C 10 min, 55°C 10 min, 50°C 20 min, 45°C 30 min, 42.5°C 15 min, 40°C 15 min, 21°C 1 min, and held at 4°C. Illumina DNA sample preparation for Tn-Seq was performed depending on the amount of gDNA collected. High gDNA amounts (>500ng) were prepared with a standard Illumina preparation method for Tn-Seq as previously described (van Opijnen & Camilli 2010). Low gDNA input amounts (*500ng) were prepared by first performing whole-genome amplification (WGA) on the gDNA sample using phi29 DNA polymerase (NEB - M0269S). 10ng of gDNA was mixed with 10μM exo-resistant primer (MCLAB - ERRP-100), 2.5mM dNTP, and 1X phi29 DNA polymerase reaction buffer, in a total volume of 26.25μl, and incubated at 95°C for 3 min and then placed on ice. Next 1XBSA (NEB), and 2 units of phi29 (0.57U/ml), were added to the reaction, and incubated for 7 hrs at 30°C, 10 min at 65°C, and held at 4°C. 10μl magnetic beads (Axygen - AxyPrep Mag PCR Clean-up Kit, MAGPCRCL50) were mixed with 30μl of freshly made PEG solution (20% PEG8000, 2.5M NaCl, 10mM Tris-base, 1mM EDTA, 0.05% tween20, pH8) and added to the 30μl sample, mixed, and incubated at room temperature for 20 min. A magnet was used to separate the bead/DNA complex from the PEG solution and the beads were washed 3 times in 200μl 70% ethanol (all magnetic bead washes were performed this way). Beads were then dried for 3 minutes at room temperature, and DNA was eluted off the beads with 12.7μl of dH_2_O. 11.49μl of phi29 amplified DNA was then added to a MmeI digestion mix (2 units NEB MmeI enzyme, 50μM SAM, 1X CutSmart Buffer) in a total volume of 20μl, and incubated for 2.5 hrs at 37°C followed by 20 min at 65°C. 1μl of alkaline phosphatase (NEB - M0290S Calf Intestinal, CIP) was added to the sample and incubated for 1 hr at 37°C. 10μl magnetic beads plus 20ul PEG solution per sample was used to wash the sample followed by elution in 14.3μl of dH_2_O. T4 DNA ligase (NEB M0202L) was used to ligate DNA adapter barcodes by adding 13.12μl DNA to 1μl of 1:5 diluted adapter, 1X T4 DNA Ligase Reaction Buffer, and 400 units T4 DNA ligase, followed by incubation at 16°C for 16 hrs, 65°C for 10 min and held at 10°C. 10μl magnetic beads plus 20μl PEG solution was used to wash the sample followed by elution in 36μl of dH_2_O. Adapter ligated DNA was then PCR amplified using Q5 high-fidelity DNA polymerase (NEB - M0491L) by adding 34μl of DNA to 1X Q5 reaction buffer, 10mM dNTPs, 0.45μM of each primer (P1-M6-GAT-MmeI; P2-ADPT-Tnseq-primer; Supplementary Table 6), 1 unit Q5 DNA polymerase, and incubated at 98°C for 30 sec, and 18-22 cycles of 98°C for 10 sec, 62°C for 30 sec, 72°C for 15 sec, followed by 72°C for 2 min and a 10°C hold. PCR products were gel purified and sequenced on an Illumina NextSeq 500 according to the manufacturers protocol. Sequence analysis was performed with a series of in-house scripts as previously described (van Opijnen et al. 2009; McCoy et al. 2017). The fitness of a single mutant (*W_i_*) is calculated by comparing the fold expansion of the mutant to the fold expansion of the population and is determined by the following equation (van Opijnen et al. 2009):

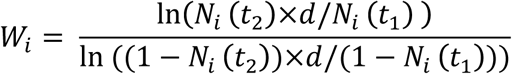

in which *N_i_(t_1_)* and *N_i_(t_2_)* are the mutant frequency at the beginning and end of the experiment respectively and *d* is the population expansion. The final average fitness and standard deviation are calculated across all insertions within a gene, and since fitness is calculated using the expansion factor of the population, *W_i_* becomes independent of time, therefore allowing comparisons between different strains and conditions across different experiments. To determine whether fitness effects are significantly different between conditions three requirements had to be fulfilled: 1) *W_i_* is calculated from at least three data points, 2) the difference in fitness between conditions has to be larger than 10% (thus *W_i_* - *W_j_* = < −0.10 or > 0.10), and 3) the difference in fitness has to be significantly different in a one sample *t*-test with Bonferroni correction for multiple testing (van Opijnen et al., 2009, 2016, van Opijnen and Camilli 2012, 2013).

### Mutant generation

Gene knockouts were constructed by replacing the entire coding sequence with a chloramphenicol or spectinomycin resistance cassette through overlap extension PCR. Construction of PCR products for gene replacement and transformation of *S. pneumoniae* were performed as described previously (van Opijnen & Camilli 2010; Iyer et al. 2005). Generated mutant strains and primers for marked deletions can be found in Supplemental Information (Supplementary Table 6,7).

### Co-culture assays

To validate genetic phenotypes associated with carbon utilization from AGP, single-gene mutants (mt) were co-cultured with their wildtype parental strain (wt) in a 1:20 ratio (mt:wt). Mutant and WT frequencies were calculated by live cell plating on blood agar plates with or without antibiotics. Fitness of the mutant was then calculated as described as previously and above (van Opijnen et al. 2009).

### Visualization of cells and droplets

Images of cells and droplets were captured with an Olympus IX83 inverted microscope. For planktonic batch culture 10μl of cells were stained with 0.5μl green-fluorescent SYTO-9 (1:10 dilution in PBS) and 0.5μl red-fluorescent propidium iodide (1:10 dilution in PBS) (Thermo Fisher Scientific - L34856). Batch culture cells were then mounted between an agar pad and coverslip for visualization. All droplet images were produced by mounting samples between coverslip spacers to prevent droplets from being compressed.

### Gene expression analysis

Immediately after culture the cells were pelleted and snap-frozen in an ethanol/dry-ice bath, followed by RNA isolation using RNeasy Mini Kit (Qiagen - 74106) according to the manufacturer's guidelines. RNA was treated to remove genomic DNA with TURBO DNA-free kit (Invitrogen - AM1907). cDNA was made from 400ng of DNA-free RNA using iScript Reverse Transcription Supermix (Bio-Rad - 1708841). Primers for quantitative real-time PCR (qRT-PCR) were designed using Primer3 software (Untergasser et al. 2012; Koressaar & Remm 2007) (Supplementary Table 6). qRT-PCR was performed with iTaq SYBR Green Supermix (Bio-Rad - 1725124) using 2μl of cDNA in a MyiQ Real-Time PCR Detection System (Bio-Rad). Each sample was measured in technical and biological triplicates and normalized to the 50S ribosomal protein SP_2204.

## Data availability

All sequence data can be found under the NCBI Sequence Read Archive accession SRP154922.

## Acknowledgements

This work was supported by R01-AI110724 and U01-AI124302 to TvO.

## Author contributions

TvO, DT and PJ conceived and worked out the idea. DW provided the initial microfluidics device and mask. PJ, DT and SW designed, produced and characterized microfluidics devices. DT performed experiments. DT and TvO performed data analyses. DT and TvO wrote the manuscript.

## Competing interests

The authors declare no competing interests.

## Acknowledgments

We would like to thank David A. Weitz and Lloyd Ung for providing, and assistance with, the initial microfluidic device. Additionally, we would like to thank Stephen Shepard at the Integrated Sciences Cleanroom at Boston College.

---

**Supplementary Figure 1:**
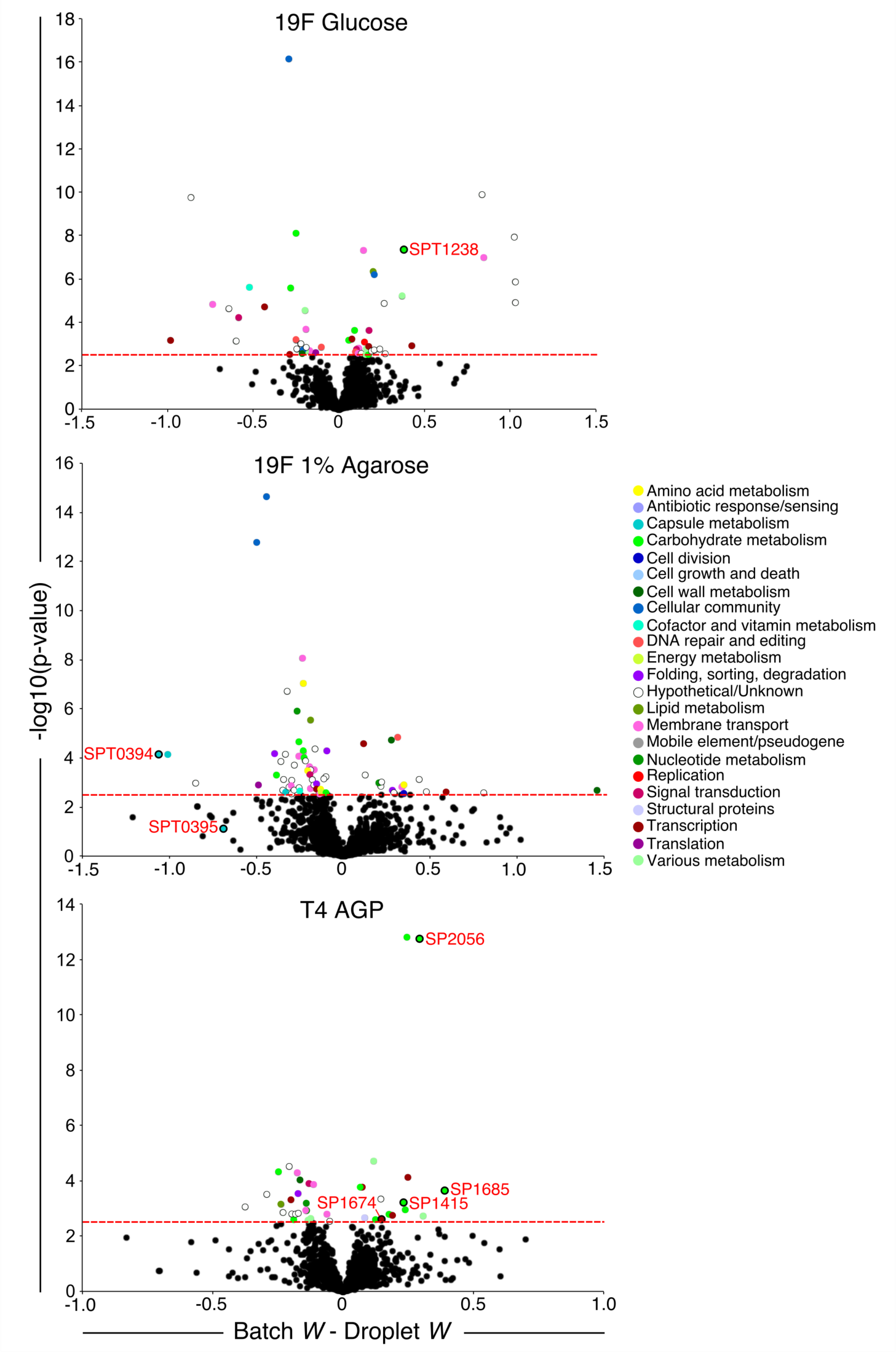
Approximately 2-5% of genes confer a significantly different fitness in droplets compared to batch culture. The functions of these significant genes span a wide range of categories including metabolism, transport, regulation and cell wall integrity. The red-dashed line indicates a conservative threshold for significance (-log(p-value)>2.5; p-value<0.003). All genes above the line are labeled with a color corresponding to a functional category represented in the figure key. The 7 genes that were validated in the study are outlined in black circles and marked with gene numbers.

**Supplementary Figure 2:**
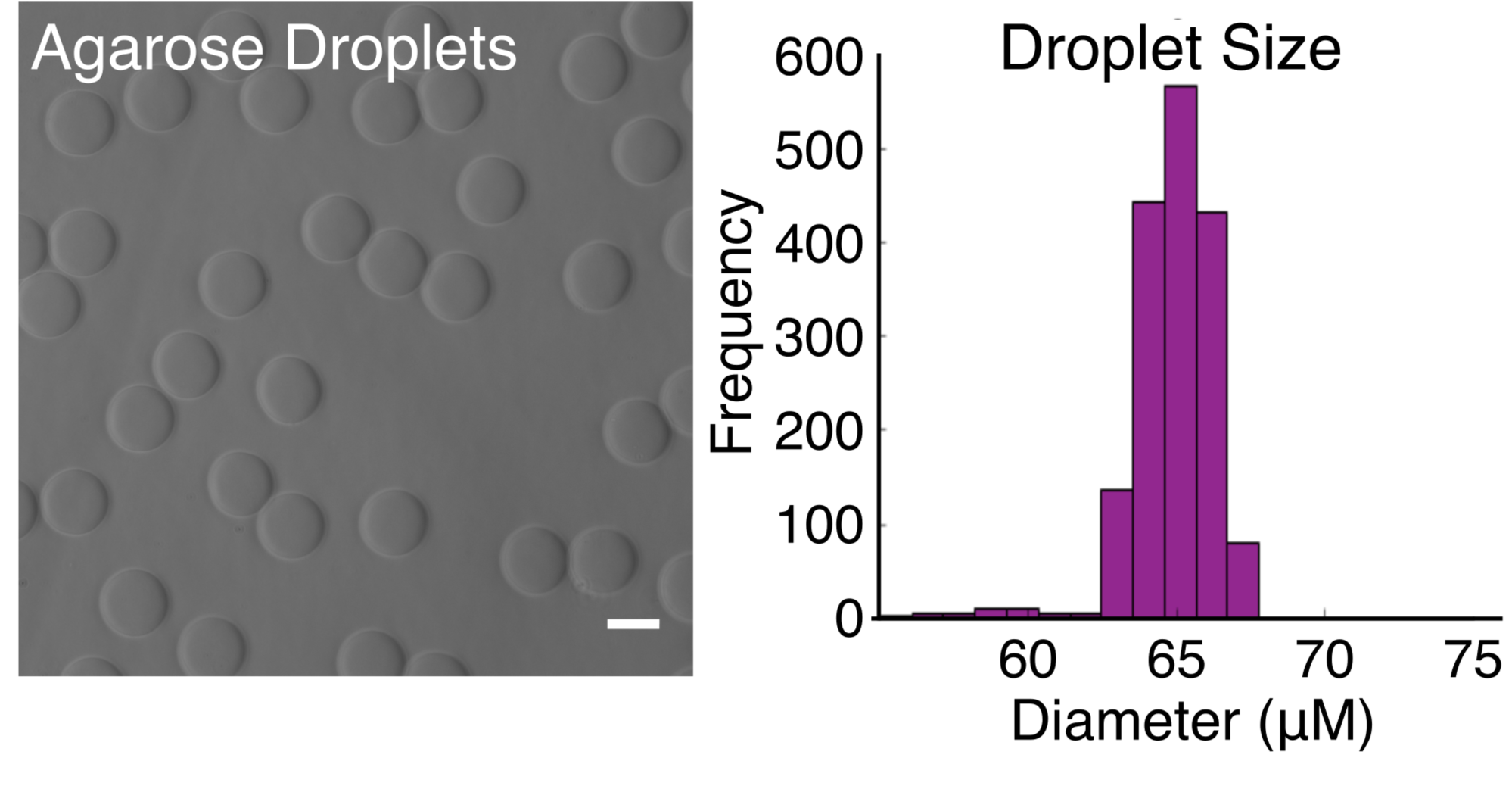
Agarose droplets with carrier oil removed. The 40μM device makes monodisperse agarose droplets that are ~65μM in diameter (~144pL volume). Scale bar is 50μM.

